# Identification of a parasitic symbiosis between respiratory metabolisms in the biogeochemical chlorine cycle

**DOI:** 10.1101/781625

**Authors:** Tyler P. Barnum, Yiwei Cheng, Kaisle A. Hill, Lauren N. Lucas, Hans K. Carlson, John D. Coates

## Abstract

A key step in the chlorine cycle is the reduction of perchlorate (ClO_4_^-^) and chlorate (ClO_3_^-^) to chloride by microbial respiratory pathways. Perchlorate-reducing bacteria and chlorate-reducing bacteria differ in that the latter cannot use perchlorate, the most oxidized chlorine compound. However, a recent study identified a bacterium with the chlorate reduction pathway dominating a community provided only perchlorate. Here we confirm a metabolic interaction between perchlorate- and chlorate-reducing bacteria and define its mechanism. Perchlorate-reducing bacteria supported the growth of chlorate-reducing bacteria to up to 90% of total cells in communities and co-cultures. Chlorate-reducing bacteria required the gene for chlorate reductase to grow in co-culture with perchlorate-reducing bacteria, demonstrating that chlorate is responsible for the interaction, not the subsequent intermediates chlorite and oxygen. Modeling of the interaction suggested that cells specialized for chlorate reduction have a competitive advantage for consuming chlorate produced from perchlorate, especially at high concentrations of perchlorate, because perchlorate and chlorate compete for a single enzyme in perchlorate-reducing cells. We conclude that perchlorate-reducing bacteria inadvertently support large populations of chlorate-reducing bacteria in a parasitic relationship through the release of the intermediate chlorate. An implication of these findings is that undetected chlorate-reducing bacteria have likely negatively impacted efforts to bioremediate perchlorate pollution for decades.

## Introduction

The chlorine cycle consists of the biological, geological, and chemical processes that interconvert organic and inorganic chlorine compounds (Atashgahi et al 2018). Chlorine oxyanions are a group of inorganic chlorine compounds of particular interest in biology due to their high reduction potentials (E^0’^ > 0.7 V) (Liebensteiner et al 2016, McCullough and Hazen 2003, Winterbourn 2008, Youngblut et al 2016b). Hypochlorite (ClO^-^) and chlorite (ClO_2_^-^) are highly reactive compounds that damage cells through oxidative chemistry (Gray et al 2013, Hofbauer et al 2016, Melnyk et al 2015), while chlorate (ClO_3_^-^) and perchlorate (ClO_4_^-^) are used as electron acceptors in respiration by some bacteria and archaea (Youngblut et al 2016b). Uniquely among chlorine oxyanions, perchlorate is chemically stable in solution, and a necessary step in the chlorine cycle is the reduction of perchlorate to chloride by microbial respiration (Coates and Achenbach 2004, Youngblut et al 2016b). Where this microbial activity is absent, geochemical reactions in the atmosphere lead to the accumulation of perchlorate and, to a lesser degree, chlorate (Kounaves et al 2010, Melnyk and Coates 2015, Youngblut et al 2016b). Both atmospheric deposition of chlorine oxyanions and microorganisms respiring chlorine oxyanions appear to be widespread (Coates et al 1999, Rajagopalan et al 2009), yet the biogeochemistry of this key part of the chlorine cycle is not well understood (Youngblut et al 2016b).

An important unresolved question is whether the microbial respiration of chlorine oxyanions in the environment is performed by individual cells or by groups of cells with different parts of the biochemical pathway (Barnum et al 2018, Clark et al 2016). Many redox metabolisms from other elemental cycles have been found to occur through pathways that are divided between different cells, including nitrate reduction (Van de Pas-Schoonen et al 2005); ammonia oxidation (Daims et al 2016, Winogradsky 1892); sulfur oxidation and reduction (Anantharaman et al 2018, Kelly et al 1997); and organic chlorine reduction (Grostern and Edwards 2006). Complete pathways might even be rare in environmental systems: a recent description of metagenome-assembled genomes from aquifer sediment found that only a minority of organisms with genes for nitrate reduction or sulfur oxidation had the complete pathway (Anantharaman et al 2016). In many cases, respiratory metabolisms have been observed to involve both cells with complete pathways and cells with partial pathways, a form of symbiosis that can range from mutualistic to antagonistic (Costa et al 2006, Dolinšek et al 2016, Hallin et al 2018, Lilja and Johnson 2016).

Chlorate reduction could be considered a partial pathway of perchlorate reduction, as the two pathways share substantial similarities (Youngblut et al 2016b). The key difference is whether or not the initial step of the pathway is catalyzed by a perchlorate reductase (Pcr), which reduces both perchlorate and chlorate, or by a chlorate reductase (Clr), which can only reduce chlorate (Figure 1A) (Wolterink et al 2003). Both metabolisms occur in the bacterial periplasm, where perchlorate and/or chlorate are reduced to chlorite, chlorite is converted to chloride and oxygen by a chlorite dismutase (Cld) (Bender et al 2002, Coates et al 1999, Hofbauer et al 2014, Van Ginkel et al 1996), and oxygen is reduced to water by one or more terminal oxidases (Clark et al 2014, Clark et al 2016, Sun 2008). Energy is conserved by the reduction of perchlorate, chlorate, and oxygen but not in the conversion of chlorite to oxygen and chloride (Figure 1A) (Rikken et al 1996). Genes for these enzymes are found together within horizontally transferred genomic DNA or plasmid DNA, typically with accessory genes for signaling and regulation, reactive chlorine stress response, protein and cofactor assembly, and genetic mobility (Clark et al 2013, Melnyk et al 2011, Melnyk and Coates 2015). Some bacteria and archaea have been experimentally observed or engineered to reduce perchlorate or chlorate to chlorite, relying on a second organism or chemical reactions to remove chlorite (Clark et al 2016, Liebensteiner et al 2013, Liebensteiner et al 2015, Martínez-Espinosa et al 2015). However, selection for perchlorate- or chlorate-reducing microorganisms from the environment has only yielded bacteria with the canonical pathways described above (Barnum et al 2018, Youngblut et al 2016b).

**Figure 1.**
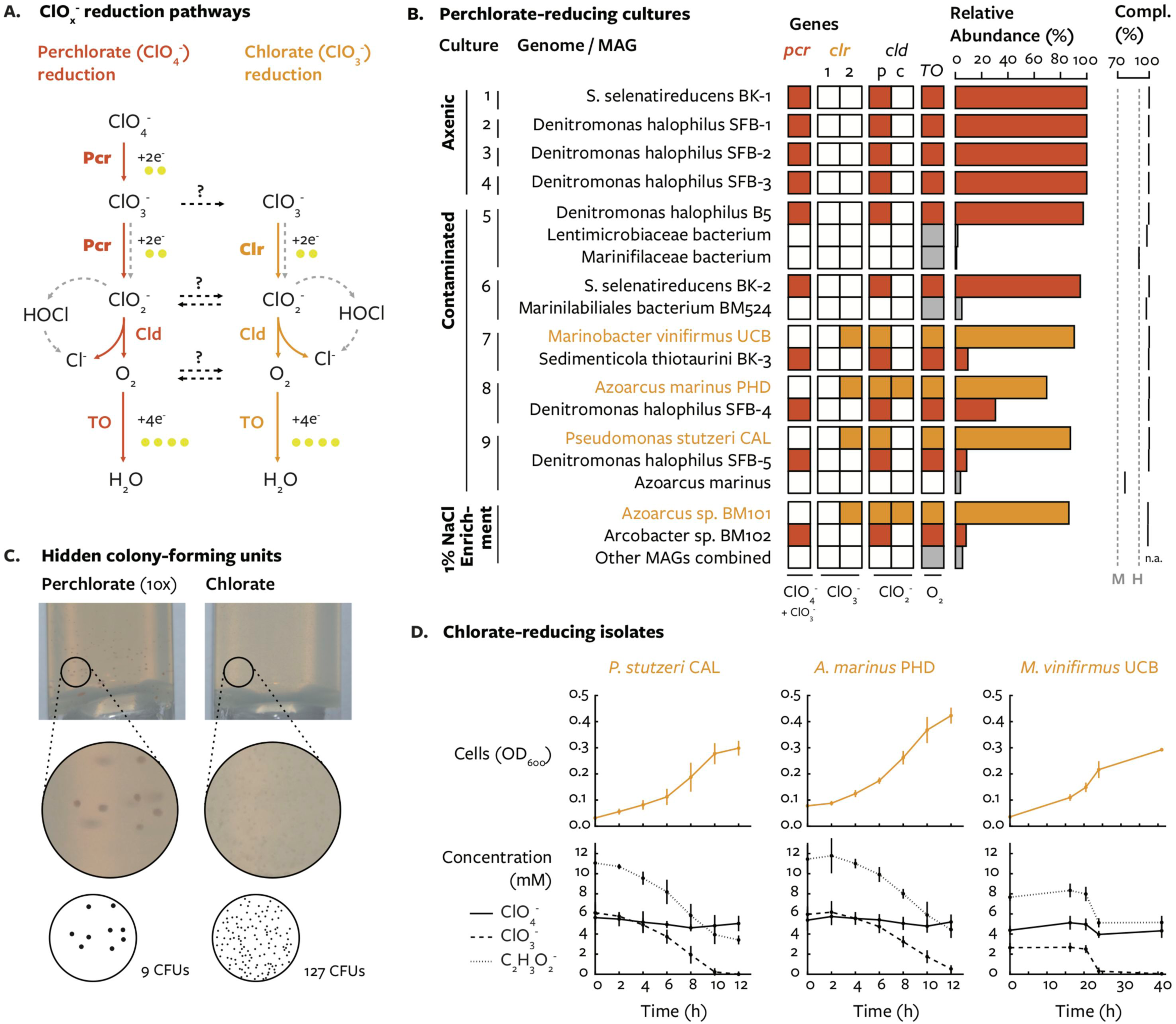
Isolation of chlorate-reducing bacteria from perchlorate-reducing cultures. (A) Pathways for the respiration of perchlorate (red) and chlorate (orange) involve the enzymes perchlorate reductase (Pcr) or chlorate reductase (Clr), chlorite dismutase (Cld), and a terminal oxidase reducing oxygen to water (TO). (B) Binning and key genes of genomes from perchlorate-reducing cultures. A previously sequenced perchlorate-reducing enrichment is included for comparison (“1% NaCl Enrichment”). Filled squares indicate gene presence. Relative abundance (%) was calculated as normalized coverage divided by total coverage for all genomes. Compl. (%) refers to percent completeness (single copy genes); dashed lines indicate medium quality (M) and high quality (H) completeness. All genomes had negligible contamination (<3%). (C) Magnified image of colonies that developed in agar media supplied perchlorate or chlorate from a co-culture of *Denitromonas halophilus* SFB-1 and *Pseudomonas stutzeri* CAL. The inoculum for perchlorate agar media was 10-times more concentrated. (D) Dissimilatory reduction of chlorate and not perchlorate by isolated chlorate-reducing bacteria.

Though the pathways for chlorine oxyanion respiration have been studied in parallel for decades (Malmqvist et al 1994, Rikken et al 1996), research on interactions between them is sparse. One set of studies explored how unusually high accumulation of chlorate by the perchlorate-reducing bacterium *Dechlorosoma sp.* HCAP-C (PCC) could support chlorate-reducing bacteria (Dudley and Nerenberg 2007, Dudley et al 2008, Salamone and Nerenberg 2006). Addition of a chlorate-reducing bacterium in co-culture with strain HCAP-C decreased the concentration of chlorate, and while models of the system suggested growth of the chlorate-reducing bacterium, the community structure *in situ* was not determined (Dudley and Nerenberg 2007, Salamone and Nerenberg 2006). Accumulation of chlorate by strain HCAP-C was proposed to occur because a single enzyme (Pcr) catalyzes two sequential reactions in the pathway (reduction of perchlorate to chlorate, and chlorate to chlorite) (Dudley et al 2008, Nerenberg et al 2006). As that trait is shared by all known perchlorate-reducing bacteria, and several perchlorate-reducing bacteria have been reported to accumulate chlorate, albeit at much lower concentrations (Cameron Thrash et al 2010, Thrash et al 2010b, Youngblut et al 2016a), it was speculated that chlorate-reducing bacteria may be a common feature of natural perchlorate-reducing communities (Nerenberg et al 2006, Salamone and Nerenberg 2006).

No subsequent research examined the possibility of interaction between chlorine oxyanion reduction pathways in communities until recently, when we observed a genome with chlorate reduction genes in a perchlorate-enriched community (Barnum et al 2018). Surprisingly, the putative chlorate-reducing population was 10-fold more abundant than the perchlorate-reducing population. Because no chlorate had been added to the cultures, the chlorate-reducing population either had unknown perchlorate reduction genes or was metabolizing intermediates of the perchlorate reduction pathway (Barnum et al 2018).

In the present study, we investigate the interaction between perchlorate-reducing bacteria and chlorate-reducing bacteria. After sequencing the genomes of perchlorate-reducing cultures obtained from estuary sediment enrichments (Carlström et al 2016), we detected contaminating bacteria that had not been completely removed during isolation. We discovered that several cultures were not predominantly perchlorate-reducing bacteria, as expected, but dominated by chlorate-reducing bacteria. We therefore used a combination of co-cultures, genetics, and modeling to confirm the interaction, define its mechanism, and explain how such a community structure could be produced. We conclude that the environmental chlorine cycle involves the interaction of a complete pathway and a partial pathway in the reduction of perchlorate to chloride.

## Materials and Methods

### Genome sequencing, assembly, binning, and annotation

Genomic DNA was extracted using a MoBio PowerSoil DNA Extraction Kit with a cell lysis protocol consisting of vortexing and heating at 70 °C for 5 min, repeated twice (MoBio Laboratories, Inc., Carlsbad, CA). DNA library preparation and DNA sequencing were performed by the Adam Arkin Laboratory or the Vincent J. Coates Genomics Sequencing Laboratory at the California Institute of Quantitative Biosciences (QB3, Berkeley, CA) using an Illumina MiSeq V2 (150PE or 250PE) and Illumina Hiseq4000 (100PE), respectively. Paired-end reads from each sample were trimmed using Sickle v. 1.33 with default parameters (Joshi and Fass 2011), error-corrected using SGA v. 0.10.15 (Simpson and Durbin 2012), and assembled using MEGAHIT v. 1.1.2 with the parameters --no-mercy and --min-count 3 (Li et al 2015). After assembly, reads were mapped back to each assembly using the Burrows-Wheeler Alignment Tool v. 0.7.10 (BWA) BWA-MEM algorithm (Li 2013). All manipulation of reads was performed on high-performance computing clusters administered by the Computational Genomics Resource Laboratory (CGRL).

Genome assemblies were screened for contamination using Anvi’o v. 3.1 (Eren et al 2015). Briefly, contigs >2,000 bp were manually binned into genomes using the hierarchical clustering generated from sequence characteristics and read coverage. When multiple genomes were present in a single assembly, contigs were binned into metagenome-assembled genomes (MAGs). Because perchlorate and chlorate respiration involve horizontally transferred genes that are subject to poor assembly, the BLAST feature in Bandage v. 0.8.0 (Wick et al 2015) was used to identify key genes and confirm their presence and absence in genomes as previously described (Barnum et al 2018). The completeness and contamination of each genome and metagenome-assembled genome was measured using CheckM (Parks et al 2015), which measures the single copy genes expected within a lineage and defines contamination as redundant genes with less than 90% amino acid identity. Structural annotation of genomes was performed using Prokka v. 1.11 (Seemann 2014), and key genes were identified using custom profile Hidden Markov models (HMMs) trained on previously confirmed proteins using HMMER v. 3.1b2 (Finn et al 2015). All reads and genome sequences are available through the NCBI Bioproject accession PRJNA387015 (Barnum et al 2018).

### Strains, media, and culture conditions

A complete set of strains and cultivation conditions are included in Supplementary Table 1. Growth medium for perchlorate-reducing cultures consisted of either a freshwater defined medium (Coates et al 1999) or a marine defined medium (Coates et al 1995) at pH 7.2 with, unless noted otherwise, 10 mM acetate as the electron donor and carbon source and 10 mM perchlorate as the electron acceptor. All media and stocks were made anaerobic by sparging with N_2_. Growth experiments were performed at 30 C in crimp-sealed tubes with an N_2_ atmosphere. Concentrations of perchlorate, chlorate, and acetate were measured using ion chromatography. Cells were quantified by optical density at 600 nm (OD600). Isolation of chlorate-reducing strains was performed by streaking twice onto aerobic solid media and confirmed by Sanger sequencing of individual colonies’ 16S rRNA genes.

### Quantification of perchlorate- and chlorate-reducing microorganisms

Primers to measure the model perchlorate-reducing bacterium *Azospira suillum* PS and model chlorate-reducing bacterium *Pseudomonas stutzeri* PDA were designed to bind variable regions of their respective small ribosomal subunit gene (16S rRNA) sequence and amplify ∼150 bp sequence. Primers to measure all chlorate-reducing bacteria were designed to bind the chlorate reductase gene (*clrA*). The *clrA* gene consists of two phylogenetic groups (here termed groups 1 and 2) with highest similarity to the alpha subunits of selenate reductase or dimethylsulfide dehydrogenase, respectively (Clark et al 2013). Specific primer selection involved identifying highly conserved sequence positions within each *clrA* group but not across closely related genes. Related genes were identified by searching the NCBI NR database with BLASTP (Camacho et al 2009). Primer-BLAST used Primer Pair Specificity to check against select genomes in the NCBI non-redundant database (Supplementary Table 2). Template DNA was quantified using qPCR with three technical replicates; a standard curve of known concentration; and SYBR qPCR Master Mix (Thermo Fisher Scientific) on a StepOnePlus qPCR machine (Applied Biosciences). Measurements were performed on four biological replicates sampled at the time of inoculation and at the last timepoint preceding stationary phase. Quantification of total extracted DNA used the Quant-iT dsDNA Assay Kit (Thermo Fisher Scientific).

The relative abundance of isolated chlorate-reducing strains in the enriched communities was determined from previous 16S rRNA gene amplicon data under Sequence Reads Archive accession SRP049563 (Carlström et al 2016). We obtained amplicon sequence variants (ASVs) to differentiate between closely related taxa by using DADA2 v.1.10 with default settings and without pooling (Callahan et al 2016, Callahan et al 2017). 16S rRNA gene sequences from representative DPRM and DCRM were compared to ASVs using BLASTN (Camacho et al 2009). Each ASV was assigned the taxonomy of the sequence with the highest percent identity above a threshold of 95% (approximately genus-level similarity). Relative abundance was calculated from the number of reads composing each ASV and the total reads per sample.

### Genetics

Genetic deletions and insertions in *Pseudomonas stutzeri* PDA were performed using protocols, strains, and plasmids from previous work (Clark et al 2016). All primers, plasmids, and strains are included in Supplementary Tables 1 and 3. Vectors were introduced into *Pseudomonas stutzeri* PDA via conjugation with *Escherichia coli* WM3064. These vectors had regions of homology allowing allelic exchange for a clean deletion, which were obtained by selection on kanamycin and counter-selection on sucrose.

### Modeling

Modeling of perchlorate and chlorate reduction used the Equilibrium Chemistry Approximation (Tang and Riley 2013), a modification of Michaelis-Menton kinetics that can account for competition between organisms for substrates and competition between substrates for an enzyme’s active site. The reaction rate for perchlorate reduction to chlorate by perchlorate-reducing bacteria is provided by Equation 1:

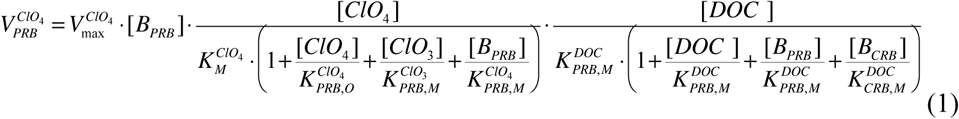

Where V (concentration time^-1^) is reaction rate and V_max_ (time^-1^) is the maximum growth rate; B, [ClO_4_^-^], [ClO_3_^-^], and [DOC] (concentration) are, respectively, the density of cells and the concentration of perchlorate, chlorate, and dissolved organic carbon (acetate). K_m_ (concentration) is the half-saturation concentration for each substrate. Together, these terms define how maximum reaction rate is limited by the concentration of perchlorate and acetate, as well as the competition of Pcr for chlorate and perchlorate.

The reaction rate for chlorate reduction was described similarly for perchlorate-reducing bacteria (Equation 2) and chlorate-reducing bacteria (Equation 3). To simplify modeling, we present chlorate reduction as one step instead of three steps (involving the intermediates chlorite and oxygen). We included the assumption that chlorate-reducing bacteria are unaffected by perchlorate.

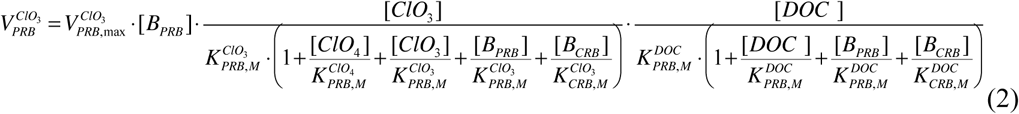

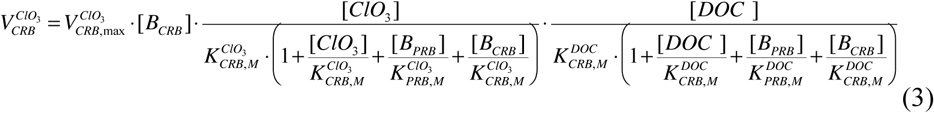

The biomass yield and stoichiometry were calculated using the framework provided by Rittmann and McCarty (Rittmann and McCarty 2001). Calculations required redox potentials and balanced half-reactions for the reduction of perchlorate to chlorate, the reduction of chlorate to chloride (via an oxygen intermediate), and the oxidation of acetate to carbon dioxide. Detailed methods and Python code are available at https://github.com/tylerbarnum/perchlorate-and-chlorate-reduction-2019. Unless otherwise noted, simulations of the model involved a theoretical case with all values equal for perchlorate-reducing bacteria and chlorate-reducing bacteria except for the ability to use perchlorate as a substrate (Table 1).

**Table 1.** Parameters for growth kinetics model. Populations were identical except for enzyme affinity for perchlorate, which for population 2 was set to be negligible (*).

| Parameter | Population 1 | Population 2 |
| --- | --- | --- |
| Km ClO<sub>4</sub><sup>-</sup> (mM) | 0.006 | 10000 * |
| Km ClO<sub>3</sub><sup>-</sup> (mM) | 0.007 | 0.007 |
| Km C<sub>2</sub>H<sub>3</sub>O<sub>2</sub><sup>-</sup> (mM) | 1 | 1 |
| Max. growth rate (h<sup>-1</sup>) | 0.5 | 0.5 |
| Death rate (h<sup>-1</sup>) | 0 | 0 |
| Type | Perchlorate reducer | Chlorate reducer |
| * Set to be negligible |  |  |

## Results

### Infiltration of perchlorate-reducing cultures by chlorate-reducing bacteria

Genomic sequencing of perchlorate-reducing cultures revealed large populations of chlorate-reducing bacteria, hinting to a metabolic interaction between perchlorate and chlorate reduction. The cultures had been obtained previously by selecting colonies from perchlorate-reducing enrichments and confirming their isolation with Sanger sequencing of the 16S ribosomal RNA gene (Carlström et al 2016). Despite appearing axenic, the nine cultures produced a total of 16 genomes after assembly and binning: four draft genomes from axenic cultures, and 11 high-quality draft metagenome-assembled genomes (MAGs) and one medium-quality draft MAG from mixed cultures (Figure 1B) (Supplementary Table 4). Every less-abundant MAG was either at low relative abundance (0.9-9.5%) or in the same taxonomic family as the most-abundant MAG, which likely caused the failure to detect contaminating strains through 16S rRNA gene sequencing. Annotation of MAGs and assembly graphs identified genes for perchlorate reduction (*pcr*, *cld*, and a terminal oxidase) in only nine genomes. Unexpectedly, while perchlorate and acetate, a non-fermentable carbon source, were the only energy substrates available in the growth medium, MAGs lacking *pcr* were the most abundant organisms in several cultures (Figure 1B). Instead, these three MAGs contained a complete chlorate reduction pathway (*clr*, *cld*, and a terminal oxidase) (Supplemental Figure 1).

The putative chlorate-reducing bacteria accounted for 69-90% of cells in the perchlorate-reducing cultures (Figure 1B), which is similar to what we previously observed in a perchlorate-enriched community (Barnum et al 2018). The dominance of putative chlorate-reducing bacteria could be visually confirmed by comparing the number of colonies that develop on anaerobic tubes containing chlorate or perchlorate as the sole terminal electron acceptor (Figure 1C). Subsequent isolation and characterization of *Marinobacter vinifirmus* UCB, *Azoarcus marinus PHD*, and *Pseudomonas stutzeri* CAL confirmed the strains to be strictly chlorate-respiring microorganisms, as no perchlorate was consumed after two weeks of incubation (data not shown) or co-metabolized during dissimilatory chlorate reduction by any strain (Figure 1D). Because these strains cannot consume perchlorate themselves, the most parsimonious explanation of the observed community structure is that a perchlorate-reducing population supported a larger chlorate-reducing population.

### Perchlorate reduction supports chlorate-reducing bacteria in simple and complex communities

The interaction between perchlorate- and chlorate-reducing bacteria was validated using defined co-cultures. Perchlorate- and chlorate-reducing strains were inoculated at equal cell densities (OD600) into anaerobic media with perchlorate as the sole electron acceptor, and the relative number of chlorate-reducing cells between inoculation and the start of stationary phase was measured using the copy number of chlorate reductase alpha subunit (*clrA*) determined by qPCR. Chlorate-reducing strains grew in every co-culture with perchlorate-reducing bacteria (Figure 2A). No growth was observed in control media that lacked an electron acceptor (Supplementary Figure 2). The fitness of the chlorate-reducing bacteria was dependent on the partner perchlorate-reducing bacterium, with *Denitromonas halophilus* SFB-1 supporting the most growth of chlorate-reducing bacteria and *Dechloromonas agitata* CKB supporting the least growth (Figure 2A). Thus, all tested perchlorate-reducing strains supported some growth of chlorate-reducing bacteria.

**Figure 2.**
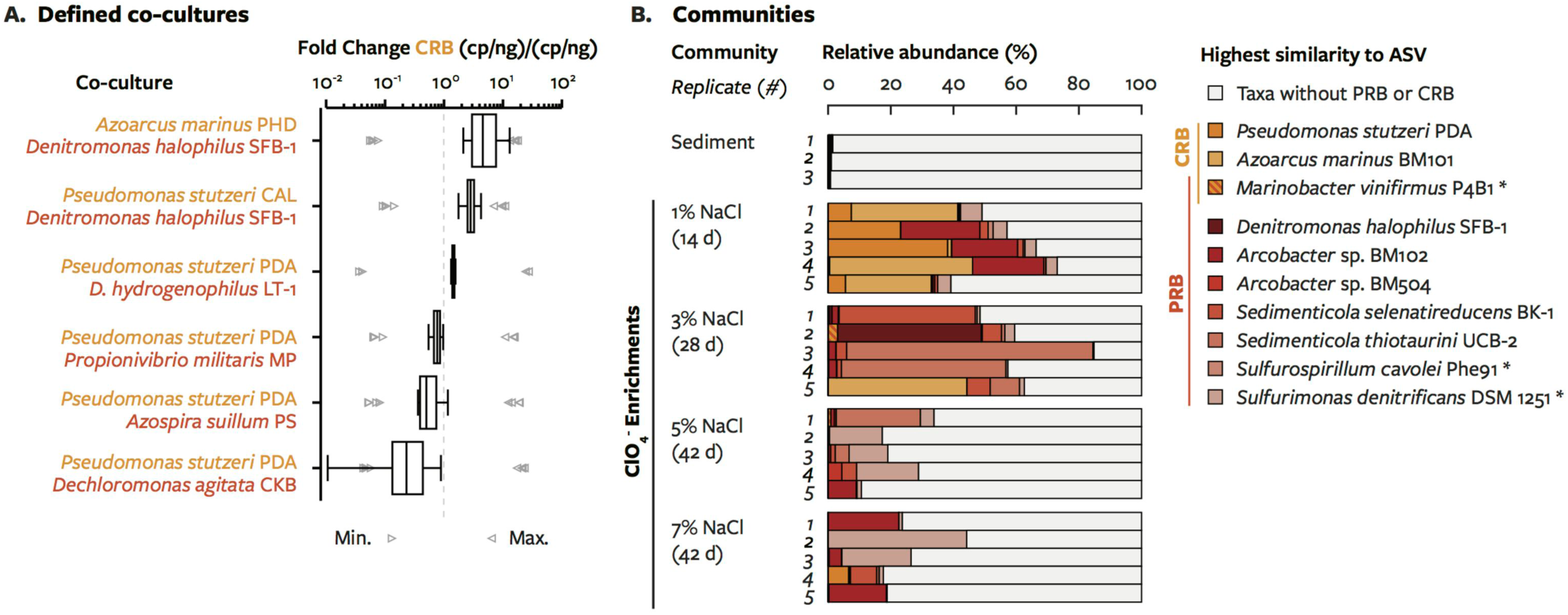
Co-cultivation of perchlorate-reducing bacteria (red, PRB) and chlorate-reducing bacteria (orange, CRB) in defined and undefined communities (A) Fold change of *clrA* in defined co-cultures between lag phase and late exponential phase batch growth. For the co-culture consisting of *P. stutzeri* PDA and *A. suillum* PS, primers for 16S rRNA genes was used. Arrows indicate the upper and lower bounds of fold change estimated from the initial and final OD600 of the co-culture. Boxplots indicate quartiles in the sample. (B) Relative abundance of 16S rRNA gene amplicon sequence variants grouped by similarity to the 16S rRNA genes of perchlorate-and chlorate-reducing taxa. *, strains most closely related to perchlorate-reducing MAGs for which 16S rRNA genes were not available.

To determine if this interaction occurs in more complex communities, we quantified the abundance of isolated strains in the original perchlorate-reducing enrichments using previously published 16S rRNA gene amplicon data (Carlström et al 2016). Indeed, amplicon sequence variants (ASVs) corresponding to isolated chlorate-reducing strains in the genera *Azoarcus* and *Pseudomonas*, which are not known to contain perchlorate-reducing species, were highly abundant (>20%) in six of ten communities at low salinity (Figure 2B). In those communities, the ASVs affiliated with chlorate reduction accounted for 23-46% of total bacteria and archaea in the community and 40-84% of putative chlorate- and perchlorate-reducing taxa. Chlorate-reducing bacteria and perchlorate-reducing bacteria were found in various combinations in communities. That many different perchlorate-reducing bacteria can support the growth of chlorate-reducing bacteria, in both co-cultures and communities, demonstrated the interaction is based not on strain-specific traits but on conserved features of the metabolic pathways involved.

### Chlorate-reducing bacteria require the perchlorate reduction intermediate chlorate

Chlorate-reducing bacteria are able to use all components of the perchlorate reduction pathway except perchlorate (Figure 1A), so we sought to determine which intermediates were responsible for the metabolic interaction. We deleted different steps of the chlorate reduction pathway in the model chlorate-reducing bacterium *Pseudomonas stutzeri* PDA (PDA). Measuring the fitness of each of these mutants in co-culture with the model perchlorate-reducing bacterium *Azospira suillum* PS (PS) would demonstrate which steps of the chlorate reduction pathway were essential for growth from perchlorate reduction intermediates. Genes encoding enzymes for reactions upstream of an exchanged intermediate are non-essential for growth, whereas genes encoding enzymes for reactions downstream of the exchanged intermediate are essential. For example, if chlorite were the exchanged intermediate, PS growing by perchlorate reduction would support growth of PDA strains lacking *clrA* but not PDA strains lacking *cld* (Figure 3A).

**Figure 3.**
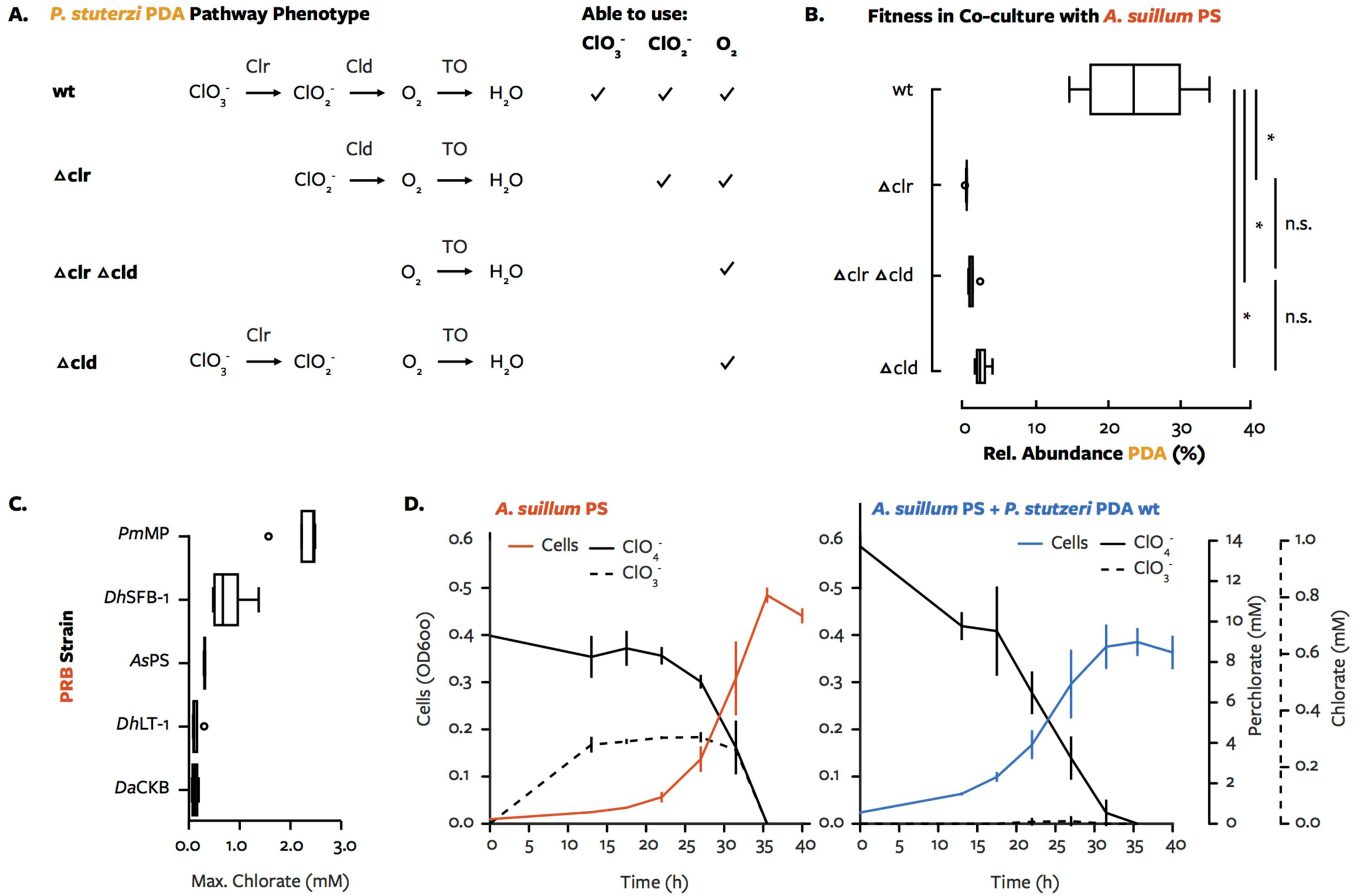
Determination of the perchlorate reduction intermediate that supports growth chlorate-reducing bacteria in defined co-cultures. (A) Genotype and phenotype (in pure culture) of chlorate reduction pathway mutants constructed in *Pseudomonas stutzeri* PDA (PDA). A chlorate reduction mutant would be unable to grow unless it can use the intermediate produced by perchlorate-reducing bacteria as a respiratory electron acceptor. (B) Fitness of chlorate reduction mutants in co-culture with *Azospira suillum* PS (PS) provided 10 mM perchlorate and 40 mM lactate, which PDA does not ferment. Relative abundance was calculated from qPCR measurements of both the PS and PDA 16S rRNA genes. *, significance of p < 0.05 (two-sided T-test); n.s., p > 0.05. Boxplots indicate quartiles in the sample with outliers as circles. (C) Maximum concentration of chlorate during dissimilatory perchlorate reduction by different strains of perchlorate-reducing bacteria (PRB) supplied 10 mM perchlorate. (D) Concentrations of perchlorate and chlorate during dissimilatory perchlorate reduction by PS or PS and PDA. Errors bars represent standard deviation of at least three replicates.

In the co-cultures inoculated with equal cell densities of PS and PDA, growth of wild type PDA was characterized by a final relative abundance of 27% (final ratio PDA/PS = 0.37) (Figure 3B). In contrast, deletion of any steps of the chlorate reduction pathway prevented growth of PDA: the final abundance of PDA deletion strains (PDA_del_/PS < 0.048) was equivalent to that expected with no growth of PDA (PDA/PS < 0.060) (Figure 3A). While all chlorate reduction genes were necessary, the particular necessity of chlorate reductase (*clrA*) demonstrated that chlorate is the only intermediate exchanged in enough quantity to support measurable growth. That is, chlorite dismutase and terminal oxidases are present in the PDA *clrA* deletion strain, yet any release of chlorite or oxygen by PS during perchlorate reduction was not sufficient to support growth of PDA. Thus, the basis of the metabolic interaction is the transfer of chlorate from perchlorate-reducing cells to chlorate-reducing cells.

Other observations supported chlorate as the exchanged intermediate. Perchlorate-reducing bacteria accumulated chlorate at concentrations between 1% and 22% mol/mol of initial perchlorate (∼10 mM) (Figure 3C), as reported previously (Thrash et al 2010a, Thrash et al 2010b). Additionally, chlorate accumulated in pure cultures of PS (<0.3 mM) but was consumed in co-cultures of PS and wild type PDA (Figure 3D). However, there was no clear relationship between the maximum concentration of chlorate that accumulated in pure cultures of perchlorate-reducing bacteria (Figure 3C) and the fitness of chlorate-reducing bacteria in co-culture (Figure 2A). Notably, kinetics differed between PS cultures with and without wild type PDA (Figure 3D, Supplemental Figure 3). When PDA was present, the maximum growth rate and maximum perchlorate reduction rate by PS decreased and the onset of growth and perchlorate reduction was earlier when compared to the pure PS culture (Figure 3D). Similar changes in growth kinetics were observed in other co-cultures (Supplemental Figure 2). The kinetics of chlorate production and consumption thus seemed to be an important factor in the interaction.

### Specificity for chlorate enables chlorate-reducing cells to exploit perchlorate-reducing cells

An understanding of the kinetics of the interaction was necessary to understand how chlorate release could produce the observed community structure. For example, how can accumulation of chlorate to only 3% of initial perchlorate concentration support chlorate-reducing bacteria at 27% of the community (Figure 3)? More generally, how can a population with the partial pathway outcompete a population with the complete pathway up to a factor of nearly ten (Figures 1-2)? To answer these questions, we used simulations of an Equilibrium Chemistry Approximation kinetics model, which included the effects of substrate competition within and between cells. We focused on the theoretical case where (1) the kinetics of chlorite and oxygen are ignored and (2) chlorate- and perchlorate-reducing populations were identical (maximum growth rate, yield, etc.) except for substrate utilization (Table 1, Equations 1-3): populations could use both perchlorate and chlorate or only chlorate. Therefore, the model’s salient features were the yields and rates from the production and consumption of chlorate, as well as the competition of perchlorate and chlorate for Pcr. In simulations with the perchlorate-reducing population alone, these parameters led to the accumulation of chlorate during perchlorate reduction (Figure 4A). Importantly, growth rate was lower while the ClO_3_^-^:ClO_4_^-^ ratio was low (Figure 4B). Chlorate influenced growth rate so strongly because chlorate reduction to chloride provided more energy (622.9 kJ/mol chlorate) than perchlorate reduction to chlorate (211.7 kJ/mol perchlorate). At low ClO_3_ :ClO_4_ ratios, the perchlorate-reducing population was less likely to reduce chlorate and more likely to reduce perchlorate (Figure 4C).

**Figure 4.**
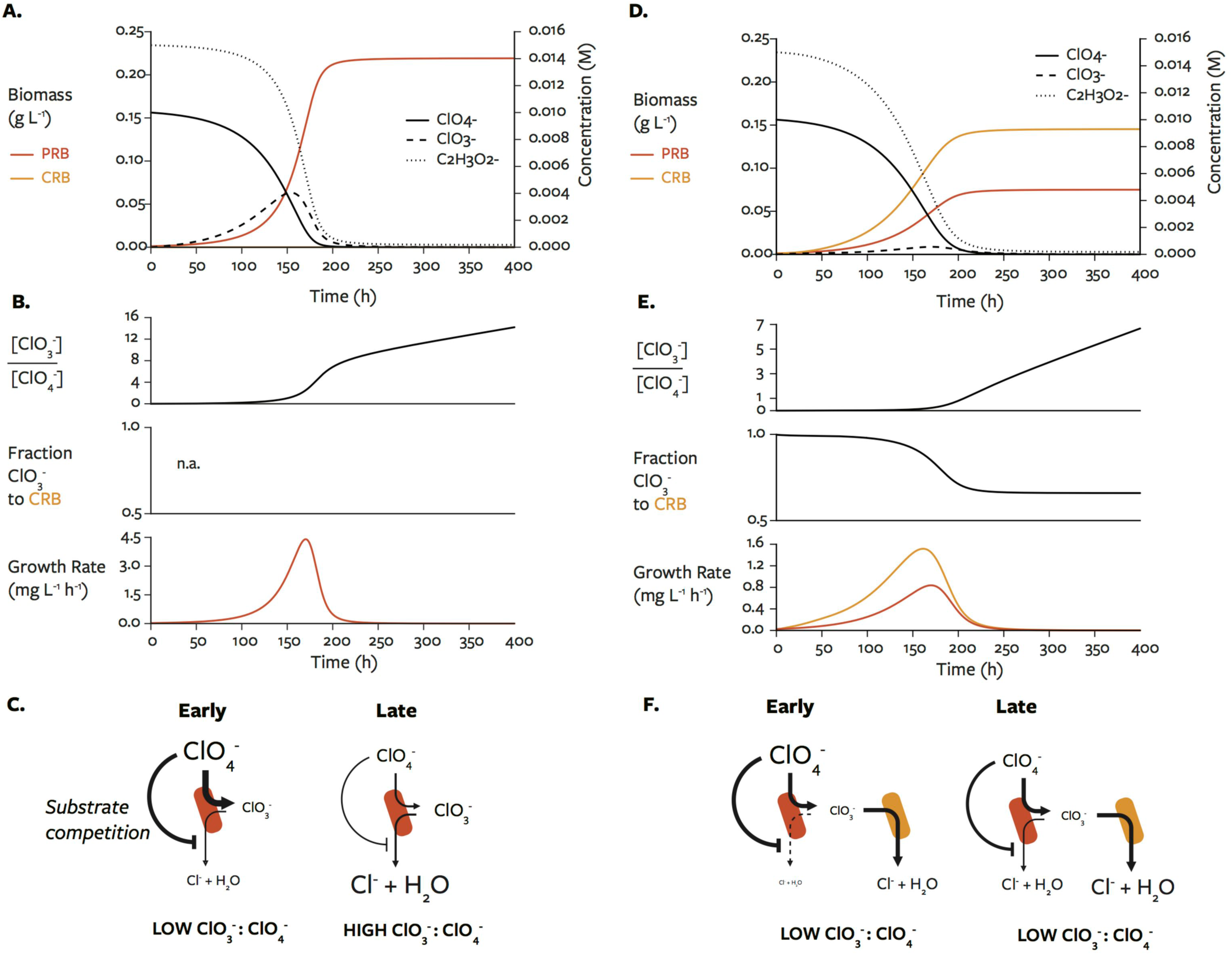
Modeling of perchlorate reduction and chlorate reduction. Simulated growth curves for perchlorate-reducing bacteria (A-C) alone and (D-F) with chlorate-reducing bacteria. [ClO_3_^-^] / [ClO_4_^-^], the ratio between chlorate concentration and perchlorate concentration; fraction ClO_3_^-^ to CRB indicates the relative amount of chlorate consumed by the chlorate-reducing population at each time step; growth rate, the change in cell concentration between each time step.

Accordingly, we hypothesized that a population that could only reduce chlorate would have a higher growth rate at low ClO_3_^-^:ClO_4_^-^ ratios than the perchlorate-reducing population. We tested this by adding the chlorate-reducing population to the simulation at equal initial concentration. The chlorate-reducing population outcompeted the perchlorate-reducing population and decreased the concentration of chlorate (Figure 4D), consistent with experimental observations. In support of our hypothesis, at low ClO_3_^-^:ClO_4_^-^ ratios the chlorate-reducing population consumed almost all of the chlorate and had a higher growth rate (about 2-fold) than the perchlorate-reducing population (Figure 4E). With chlorate-reducing cells present, the consumption of chlorate delayed the increase of the ClO_3_^-^:ClO_4_^-^ ratio (Figures 4B and 4E). Thus, in this simple theoretical case, chlorate-reducing cells had a growth advantage because they, unlike perchlorate-reducing cells, could consume chlorate at high perchlorate concentrations (Figure 4F). Additionally, the consumption of chlorate by chlorate-reducing bacteria created a positive feedback by maintaining a low ClO_3_^-^:ClO_4_^-^ ratio (Figures 4C and 4F).

We used additional simulations to observe how initial conditions affect the interaction. Varying the initial ratio of chlorate-reducing cells to perchlorate-reducing cells did not alter the ecological success or the fraction of chlorate acquired by the chlorate-reducing population (Supplemental Figure 4A-B); Chlorate-reducing cells ultimately dominated by acquiring a large percent of chlorate unless initially outnumbered 100-fold (Supplemental Figure 4C). Varying perchlorate concentration, however, did alter the success of chlorate-reducing cells (Supplemental Figure 4D-F). Even when chlorate-reducing cells outnumbered perchlorate-reducing cells, the perchlorate-reducers consumed nearly all available chlorate except at perchlorate concentrations above ∼1 mM (Supplemental Figure 4E). Also, varying the affinity of different populations for perchlorate or chlorate altered the ecological success of chlorate-reducing cells (Supplemental Figure 5). While not necessarily predictive of behavior in the environment or over different temporal and spatial scales, these simulations provide an intuitive description of the interaction: chlorate-reducing cells exploit a niche made available by differences in enzyme kinetics and substrates.

## Discussion

This study confirms and further interrogates the interaction between perchlorate reduction and chlorate reduction. Here we clearly demonstrate that bacteria with the perchlorate reduction pathway supported – and could be outcompeted by – bacteria with the chlorate reduction pathway. The interaction between perchlorate and chlorate reduction occurred in both controlled (i.e. co-cultures) and uncontrolled systems (i.e. enrichment and isolation) and in both freshwater and marine conditions. The basis of the interaction was the exchange of chlorate from perchlorate-reducing cells to chlorate-reducing cells. Chlorate was available for consumption likely due to competition of perchlorate and chlorate for a single enzyme in the periplasm of perchlorate-reducing cells (Dudley et al 2008). Simulations showed that the chlorate-reducing cells are successful because chlorate can be reduced even at a low ClO_3_^-^:ClO_4_^-^ ratio, a state that chlorate consumption perpetuates. In summary, chlorate-reducing bacteria were a common feature of perchlorate reduction and had a large effect on the structure and function of perchlorate-reducing communities.

The basis of the interaction alters our understanding of the chlorine cycle. Perchlorate reduction involves the combined activity of perchlorate-reducing microorganisms, chlorate-reducing microorganisms, and any chemical reduction of their intermediates (Figure 5). A role for chlorite-consuming or oxygen-consuming partial pathways in perchlorate reduction was not observed here (Figure 3), and an interaction based on the exchange of chlorite has been engineered (Clark et al 2016) but not yet observed in nature. This is likely due to the high activity (*k*_cat_⁄K_M_) of chlorite dismutase (10^6^-10^8^ M^−1^ s^-1^) relative to perchlorate reductase (∼10^5^ M^−1^ s^-1^) (Dubois 2014, Youngblut et al 2016a). Because chlorate is less reactive than chlorite, cells that inadvertently reduce chlorate to chlorite would experience greater reactive chlorine stress; chlorite dismutase (Cld) can detoxify chlorite produced in this manner (Celis et al 2015). Additionally, the exchange of chlorate is less constrained by the reducing state of the environment than the exchange of chlorite. But chlorate does react with common environmental reductants such as reduced iron minerals (Brundrett et al 2019, Engelbrektson et al 2014), and the reactivity of chlorate with iron increases with salinity (Brundrett et al 2019), which may contribute to the lower frequency of chlorate-reducing bacteria in higher-salinity perchlorate-reducing enrichments (Figure 2B). The concentrations of reductants, chlorate, and perchlorate may all influence the relative contribution of perchlorate-reducing microorganisms and chlorate-reducing microorganisms to chlorine oxyanion respiration.

**Figure 5.**
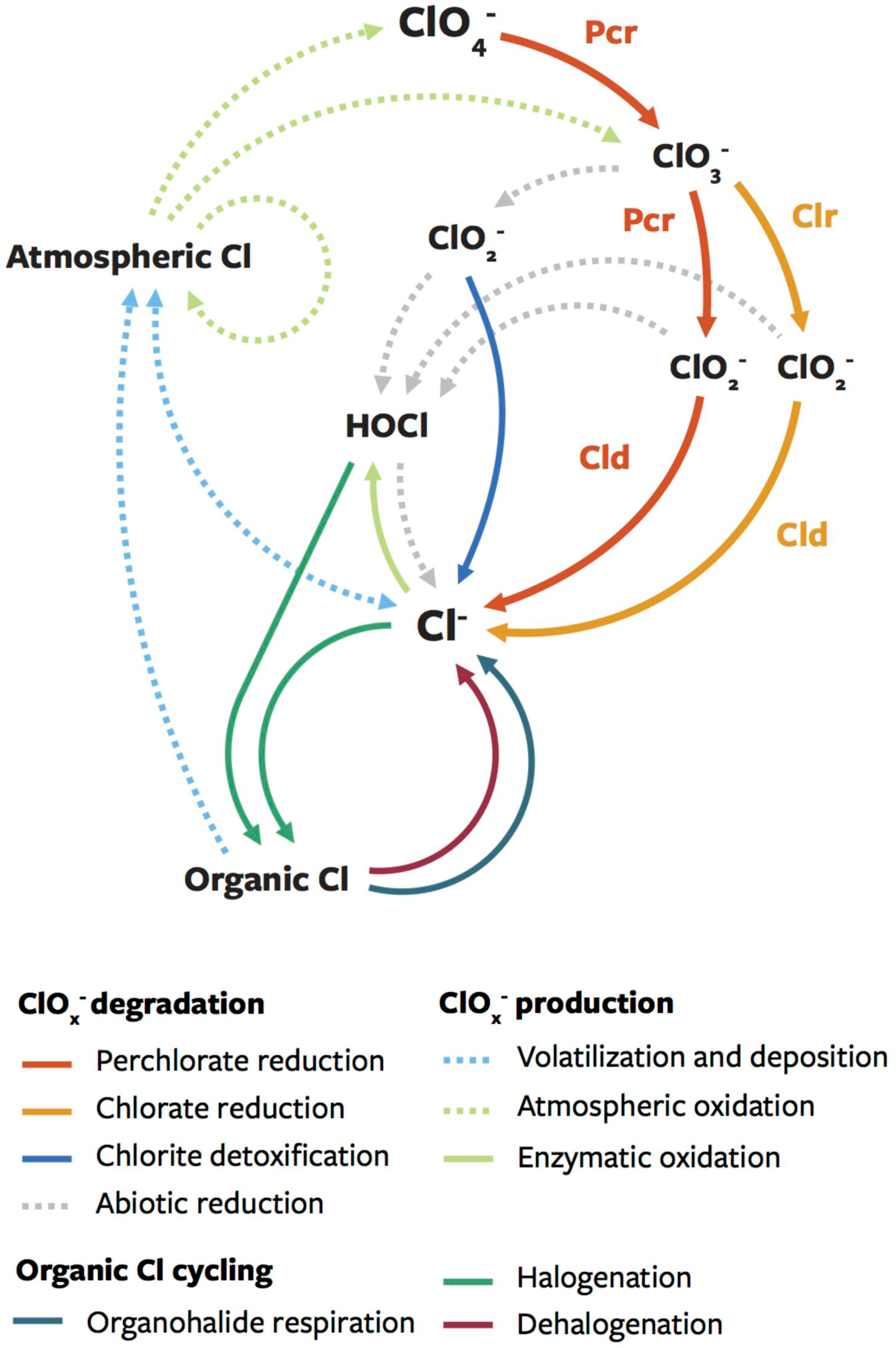
Model for the production and degradation of chlorine oxyanions. The perchlorate reduction pathway (red) accumulates chlorate, which can react with reductants and generate reactive chlorine species (gray) or be consumed by the chlorate reduction pathway (orange). We did not find evidence for the release of chlorite and oxygen by the perchlorate and chlorate reduction pathways, though both chemicals can react with any reductants in the periplasm. Perchlorate and chlorate reduction remove the products of atmospheric oxidation of chlorine (dashed yellow). Co-metabolic or inadvertent enzyme activities are not shown.

Interactions like that described here, where low accumulation of an intermediate supports large populations with a partial respiratory pathway, may be common across elemental cycles. Some evidence exists for the importance of these interactions in denitrification, for example. *Pseudomonas* strain G9, which contains a complete denitrification pathway producing inhibitory concentrations of nitrite, could grow only in co-culture with *Alcaligenes faecalis* strain TUD, which only reduces nitrite to dinitrogen, and the two strains were found at steady state at approximately equal cell densities (Van de Pas-Schoonen et al 2005). Nitrite accumulation caused by inter-enzyme competition in *Pseudomonas stutzeri* strain A1501 was decreased by dividing nitrite production and consumption between different strains (Lilja and Johnson 2016). In denitrifying communities, then, it may be beneficial for some organisms to *lack* steps in the denitrification pathway. This was the case for a pooled transposon mutant library of *Azospira suillum* PS, where mutants with insertions in nitrite reductase, which is deleterious in pure culture, outcompeted cells with intact denitrification pathways (Melnyk et al 2015). Much remains to be learned about community structure impacts resulting from pathway distribution across different populations.

The importance of studying metabolic interactions in biogeochemical transformations is to learn how such interactions influence concentrations and rates. Previous studies that added chlorate-reducing bacteria to cultures of the perchlorate-reducing bacterium HCAP-C, which accumulates far more chlorate than the typically observed (Cameron Thrash et al 2010, Thrash et al 2010b, Youngblut et al 2016a), had conflicting results where chlorate-reducing bacteria either slightly accelerated (Salamone and Nerenberg 2006) or substantially decelerated the rate of perchlorate reduction (Dudley and Nerenberg 2007). We observed that adding chlorate-reducing bacteria to cultures of the model perchlorate-reducing bacterium *Azospira suillum* PS decreased the concentration of chlorate, maximum growth rate, and maximum perchlorate reduction rate (Figure 3D). Similar effects on growth rate were observed with perchlorate-reducing bacteria that accumulated varying concentrations of chlorate (Supplemental Figure 2, Figure 3C), and we directly demonstrated success of chlorate-reducing populations at the expense of perchlorate-reducing populations (Figure 2). Because chlorate reduction appears to substantially influence concentrations and rates during perchlorate reduction, chlorate-reducing bacteria could affect efforts to bioremediate perchlorate. For example, a description of perchlorate-reducing bioreactors with gene-centric metagenomics identified *Azoarcus* and *Pseudomonas* as among the most abundant genera, yet the *Azoarcus* isolate did not reduce perchlorate (Stepanov et al 2014). Our results predict that those organisms are chlorate-reducing bacteria that persisted in the bioreactors for over 10 years (Stepanov et al 2014). Understanding how this metabolic interaction affects perchlorate reduction kinetics in different systems, and how it could be controlled, may be a promising line of future research.

A close interaction between metabolisms also has evolutionary implications, as co-occurrence can influence gene evolution and exchange. For example, in nitrifying microorganisms, niche differentiation led to high affinity and low affinity ammonia-oxidizing enzymes that function best at different pH (Martens-Habbena et al 2009) and to the exchange of ammonia monooxygenase to nitrite-oxidizing bacteria (Daims et al 2015). Not enough chlorate reductases and perchlorate reductases have been evaluated to draw general conclusions from their substrate affinities and catalytic rates. However, several chlorate-reducing bacteria contain genes or gene fragments of perchlorate reductase components (*napC* and *pcrD*) adjacent to the chlorite dismutase (*cld*), and these genes were most likely acquired from perchlorate-reducing bacteria (Clark et al 2013). Because environmental reduction of perchlorate or chlorate will likely involve both perchlorate-reducing microorganisms and chlorate-reducing microorganisms, a history of gene exchange between the two metabolisms is unsurprising.

## Conclusions

Perchlorate reduction supports chlorate reduction through the release of the intermediate chlorate. The fundamental cause of the interaction is that the perchlorate reductase enzyme catalyzes both perchlorate reduction to chlorate and chlorate reduction to chlorite – therefore chlorate competes with perchlorate for perchlorate reductase, limiting subsequent steps of the perchlorate reduction pathway. Chlorate reduction, despite being a partial pathway, is ecologically successful because it can consume chlorate unabated and, in doing so, exacerbates the imbalance between perchlorate and chlorate. As for several other respiratory metabolisms, the respiration of chlorine oxyanions in the environment should be expected to involve cells performing complete and partial respiratory pathways. These findings have clear implications for understanding the evolution and the kinetics of chlorine oxyanion reduction.

## Acknowledgements

Financial support was provided through a grant from the Energy Biosciences Institute EBI-BP program to JDC and through the NSF Graduate Research Fellowship Program to TPB. We thank Kelly Whetmore and the Adam Arkin Laboratory for performing library preparation and sequencing.

## Contributions

JDC guided the research. LNL and TPB isolated strains. LNL extracted DNA for sequencing. TPB assembled and analyzed genomes, designed primers, and performed amplicon sequence variant analysis. TPB and KAH performed all experiments and measurements. YC developed the model. YC and TPB performed and interpreted modeling simulations. TPB wrote the manuscript and created the figures with guidance from JDC. All authors contributed to data analysis, reviewed the manuscript, and approved of its publication.

---

Supplemental Figure 1. Visualization of assemblies and binning using Bandage to verify binning of key genes. Only mixed cultures with perchlorate-reducing bacteria and chlorate-reducing bacteria are shown: cultures (A) “UCB,” (B) “CAL,” and (C) “PHD.” The de Bruijn graph assembly is visualized by displaying contigs (lines), and connections between contigs that could not be resolved during assembly. Thickness of lines indicates sequencing depth. In each assembly, contigs with the same color were found in the same bin. Bins containing contigs with chlorate reduction genes (*clr, cld*) are colored orange, bins containing contigs with perchlorate reduction genes (*pcr, cld*) are colored red, and bins without either set of genes are other colors. Arrows indicate the contig(s) with key genes. Mean sequencing depth for those contigs and each genome bin are indicated in parentheses.

Supplementary Figure 2. Growth phenotypes of the different combinations of perchlorate- and chlorate-reducing bacteria. Individual replicates are shown due to interesting variation. Blue, co-cultures with 10 mM perchlorate; red, perchlorate-reducing bacteria with 10 mM perchlorate; solid orange, chlorate-reducing bacteria with 10 mM chlorate; and dashed orange, chlorate-reducing bacteria with no electron acceptor.

Supplemental Figure 3. Growth curves of chlorate reduction pathway mutants in *Pseudomonas stutzeri* PDA (PDA) in co-culture with *Azospira suillum* PS (PS). Errors bars represent standard deviation of four replicates.

Supplementary Figure 4. Simulations of the kinetics-based model that varied the initial concentrations of perchlorate-reducing bacteria and chlorate-reducing bacteria (A, B, C) or perchlorate-reducing bacteria and perchlorate (D, E, F). For each simulation, the final ratio of the populations (A, C) and the total percent of chlorate consumed by the chlorate-reducing population (B, E) were determined after 1000 hours with 1-hour time steps. (C, F) depict the relationship between the two measurements. The default conditions used in the main text are highlighted in white: 10^-5^ M (∼0.001 g/L) cells and 10 mM perchlorate.

Supplementary Figure 5. Simulations of the kinetics-based model that measured the final ratio of chlorate-reducing bacteria to perchlorate-reducing bacteria after varying (A) the affinity of perchlorate-reducing bacteria for chlorate and perchlorate and (B) the affinity of chlorate-reducing bacteria and perchlorate-reducing bacteria for chlorate. The default conditions used in the main text are highlighted in white: 6 μM Km for chlorate (for both populations) and 6 μM Km for perchlorate (for perchlorate-reducing bacteria only).

