## Supplementary figures and images for "Identification of a parasitic symbiosis between respiratory metabolisms in the biogeochemical chlorine cycle"

### Supplementary Figures 1-5

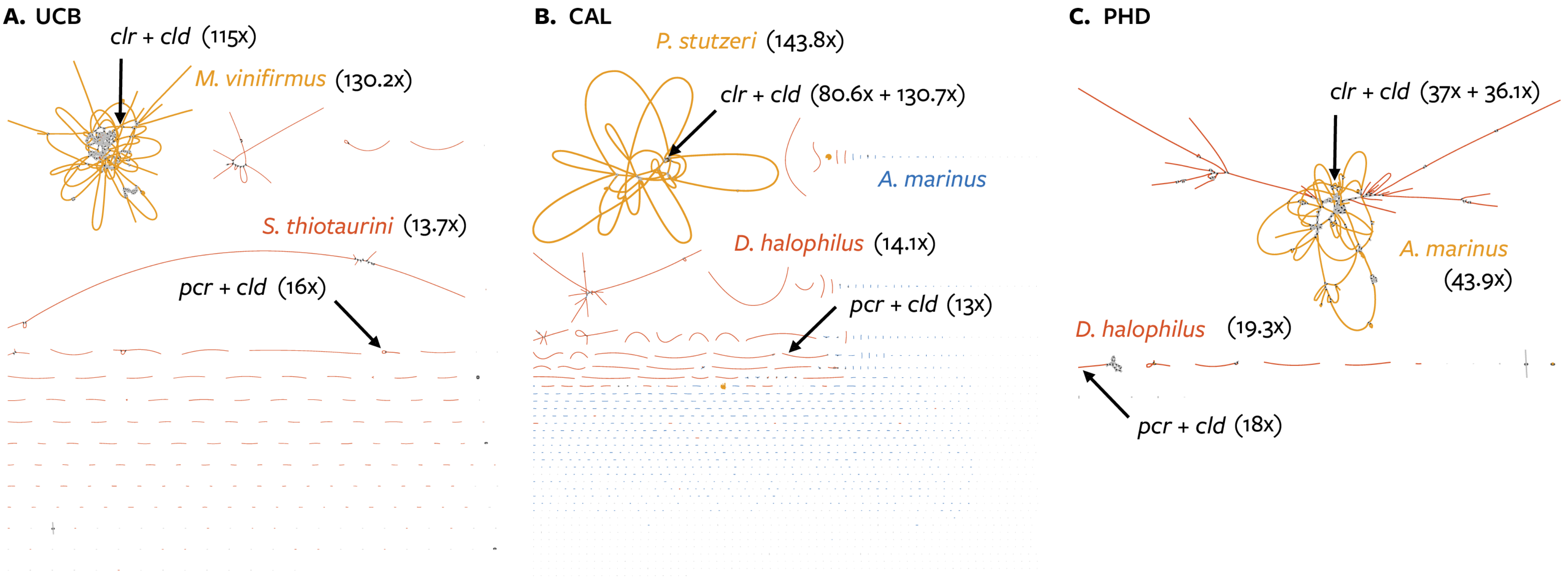

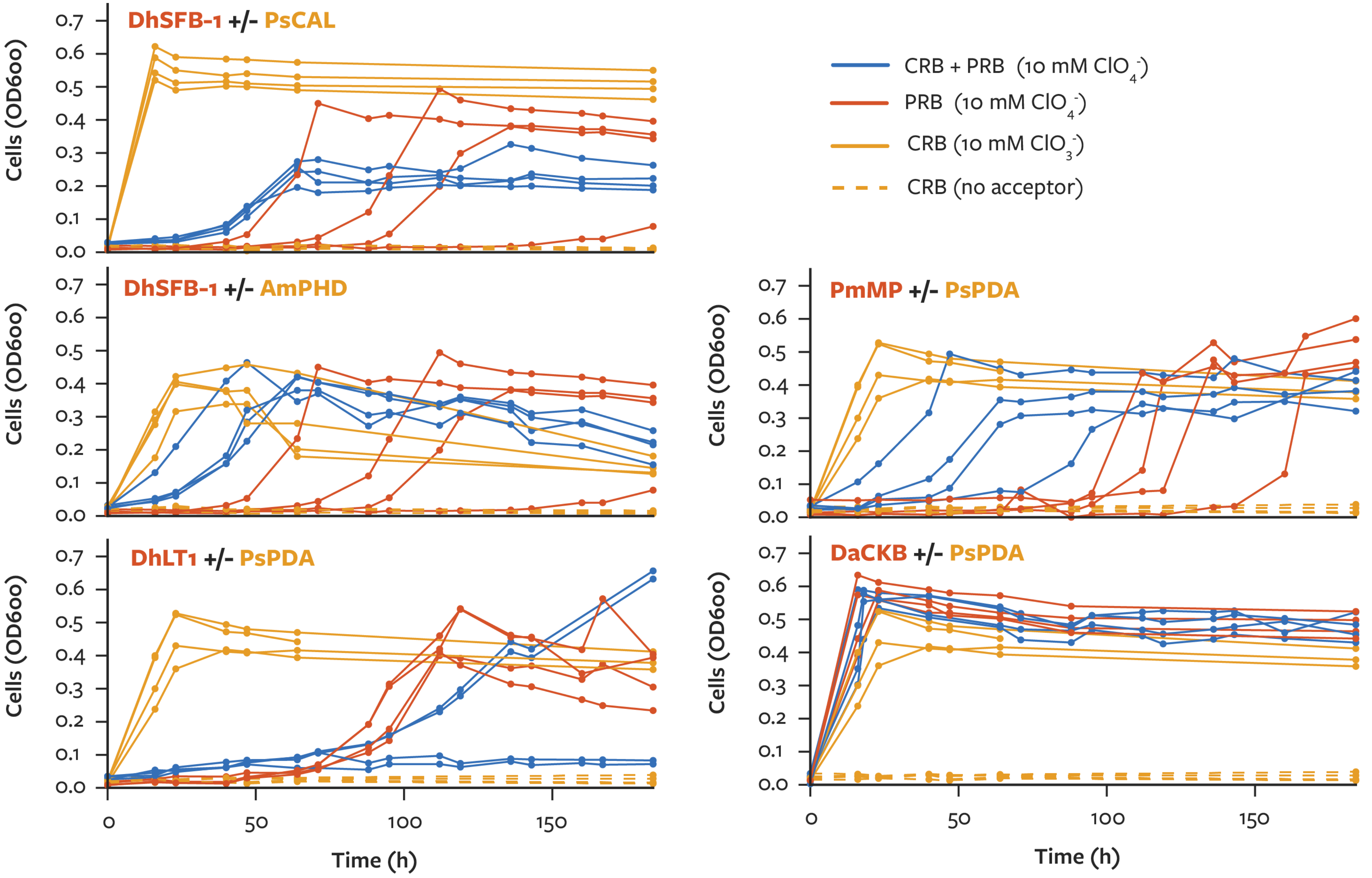

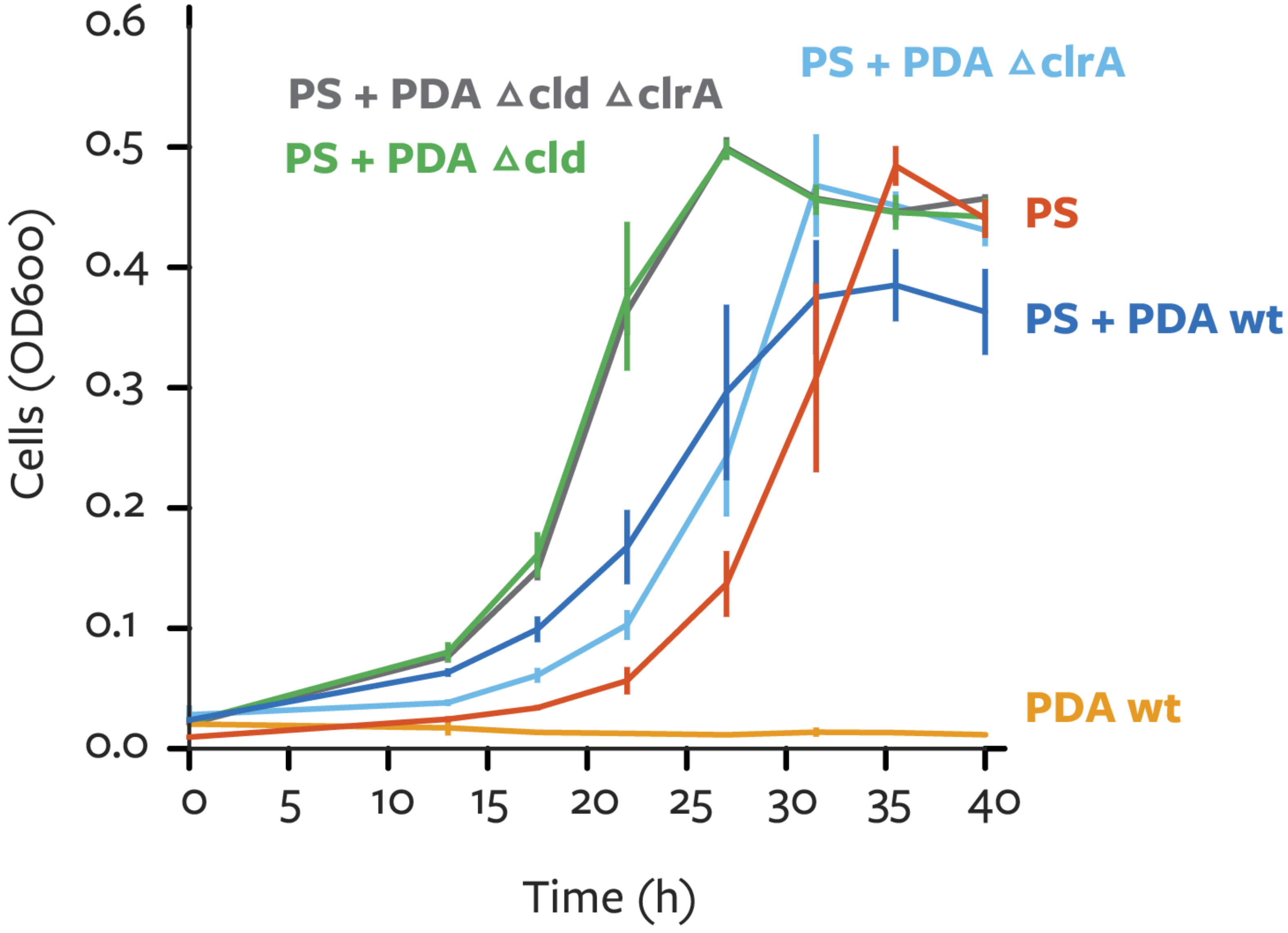

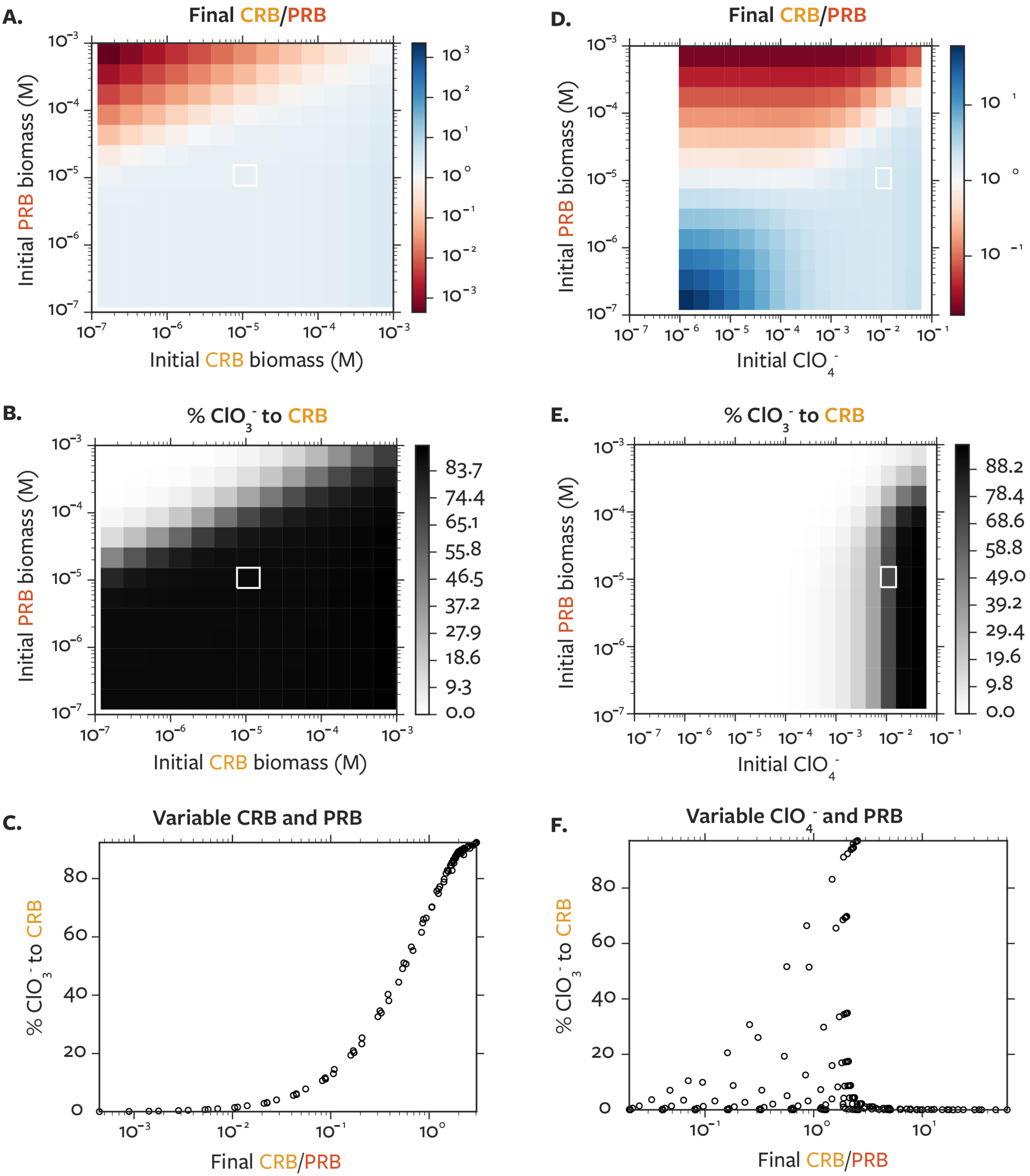

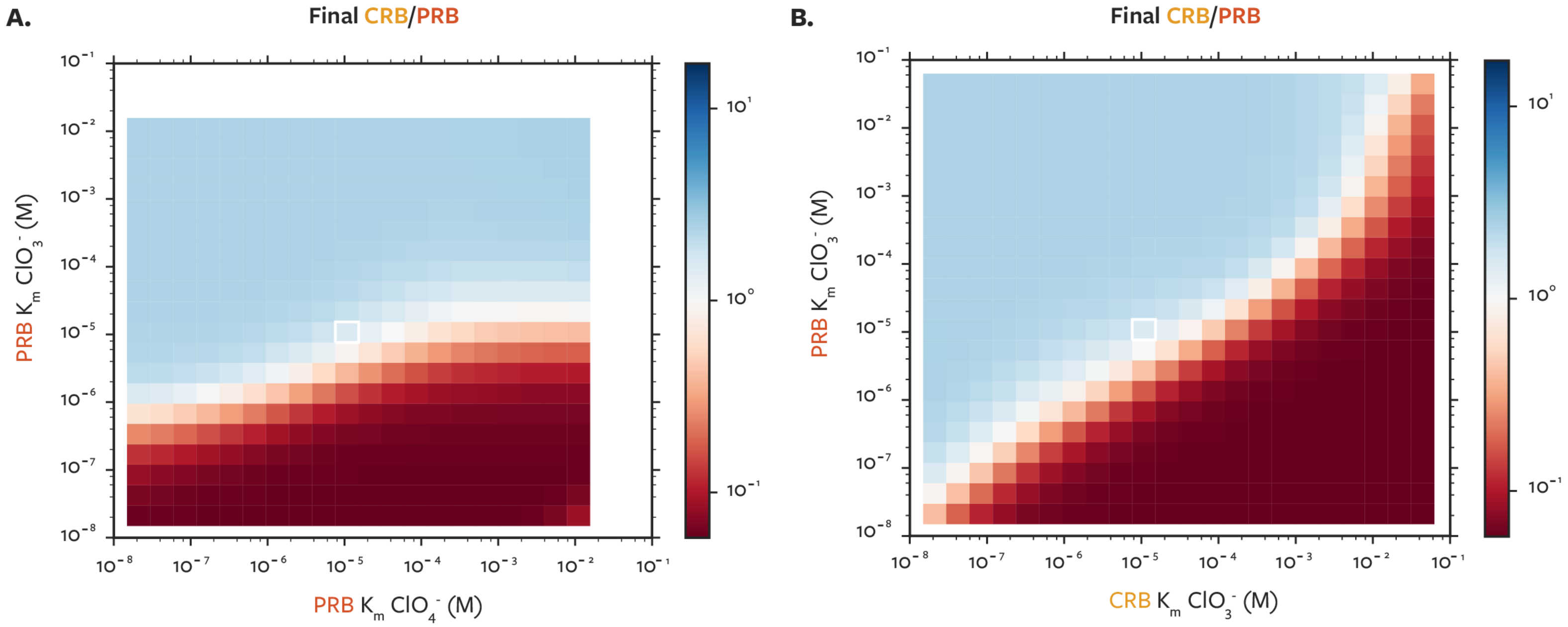
